# Residual Multi-Modal Learning for Pan-Breast-Cancer Drug Response Prediction

**DOI:** 10.64898/2026.07.03.736239

**Authors:** Beibei (Bay) Huang, Lance Tasaka, Jiaheng Li, Tanvir Islam, Shuxing Zhang

## Abstract

Predicting drug sensitivity across diverse cancer cell lines remains a fundamental challenge in precision oncology, particularly for data-scarce cell lines where per-cell-line models overfit and lookup-table approaches cannot generalise to unseen biological contexts. We present **DL4DR**, a Two-Tower Residual Late Fusion deep learning model that addresses this challenge through content-based, identity-free genomic conditioning. The Cell Line Tower encodes each cell line as a 3 × 139 × 139 genomic image - encoding gene expression, mutation severity, and copy-number variation as RGB channels - using a convolutional encoder that maps directly from biological content, never from a cell line ID. The Compound Tower combines three complementary molecular representations: D-MPNN graph message passing, ORNN octave convolutional image features, and an ECFP hard-memorization head that preserves activity-cliff resolution. Predictions are composed as a residual sum: *f* = *f*_hard_ + *λ*(**z**_*C*_) · *f*_residual_, where the learned gate *λ* modulates how much interaction signal supplements the memorization baseline.

Evaluated across **51 breast cancer cell lines** (136,342 records), Residual Fusion outperforms the ECFP-Only baseline in 48/51 cell lines (94.1%), with Δ*R*^2^ > 0.02 in 26/51 (51.0%). On the leave-cell-line-out split - the decisive test of genomic generalisation - the mean Δ*R*^2^ = +0.016 across all 51 lines demonstrates that the genomic encoder learns transferable biological signal beyond cell line identity. External validation on 601 cell lines across 27 cancer tissue types (CellTiter-Glo dataset; 0 cell line overlap with training) achieves median *R*^2^ = 0.627, within the range of the internal random-split performance (*R*^2^ = 0.61-0.69), confirming pan-cancer generalisation. GradCAM interpretability on the Cell Line Tower recovers TP53 among the top-five cross-cell-line genomic activators (5/51 cell lines) alongside several uncharacterised candidate genes (e.g. FSIP2, 6/51) - without any prior pathway annotation - providing partial biological validation of the learned representation, while also indicating that a substantial share of the encoder’s top-ranked signal corresponds to genes with no current annotation as breast cancer drivers. Code and data are available at https://github.com/bayjuan5/DL4DR.

## 1 Introduction

Accurately predicting how individual cancer cell lines respond to chemical compounds is a central task in computational oncology [Chaudhari et al., 2017, 2020]. Large-scale pharmacogenomic datasets — notably the Genomics of Drug Sensitivity in Cancer (GDSC) [Garnett et al., 2012, Yang et al., 2013] and Cancer Cell Line Encyclopedia (CCLE) [Barretina et al., 2012] — have provided compound-cell-line IC_50_ matrices with tens of thousands of measurements, enabling machine learning approaches that jointly model molecular structure and genomic context [Yang et al., 2019, Nguyen et al., 2022, Saeed et al., 2025].

Two dominant paradigms have emerged. *Fingerprint-based approaches* (ECFP + MLP) exploit the memorization power of extended-connectivity fingerprints to resolve activity cliffs — abrupt potency changes caused by minor structural modifications — but cannot generalize to structurally novel compounds [Rogers and Hahn, 2010, Stumpfe and Bajorath, 2012, van Tilborg et al., 2022]. *Graph neural networks* (GCN, D-MPNN) [Kipf and Welling, 2017, Gilmer et al., 2017] overcome this limitation through learnable molecular representations, but suffer from over-smoothing as depth increases, causing node representations to converge and erasing the localized structural information that distinguishes activity cliffs [Dablain et al., 2024].

Cell line representation presents a complementary challenge. Most existing models parameterize cell lines using a lookup table: a learned embedding vector indexed by cell line identity. This approach is fundamentally limited for generalization: any cell line absent from training has no entry in the table, and predictions for unseen biological contexts are impossible. Given that practical drug repurposing requires predictions for novel patient-derived cell lines or organoids, this limitation is not merely theoretical.

Recent work has made significant progress on multi-modal fusion architectures for drug response prediction. DeepDTF [Zhao et al., 2026] proposes a dual-branch Transformer framework that encodes multi-omics profiles via a CNN-Transformer and drug structures via a GNN–Transformer, fused through a cross-modal attention module, achieving strong performance on cold-start cell line evaluation (*R*^2^ = 0.875, AUC = 0.987 with full multi-omics). However, Deep–DTF and related approaches [Nguyen et al., 2022, Saeed et al., 2025] still represent cell lines as identity-indexed feature vectors: the omics profile is used as input, but the model implicitly assumes the cell line is present at inference time through the data split structure. DL4DR takes a fundamentally different approach: the Cell Line Tower encodes genomic state as a content-based image with no identity dimension, enabling *true* zero-shot generalization to cell lines never encountered during training. This distinction is validated by our external evaluation on 601 cell lines spanning 27 cancer tissue types with zero overlap with the training set.

We address both challenges simultaneously with **DL4DR**, a Two-Tower Residual Late Fusion architecture:

1. **Compound Tower:** combines ECFP hard memorization with D-MPNN graph learning and ORNN octave image features via a residual fusion that structurally prevents the learnable branch from overwriting memorized activity-cliff knowledge.
2. **Cell Line Tower:** encodes genomic state as a 3 *×* 139 *×* 139 RGB image using a CNN that maps from biological content — gene expression, mutation severity — without any cell line identity dimension. Any cell line with a genomic profile can be encoded at inference time, including ones never seen during training.
3. **Residual Fusion:** decomposes predictions into additive main effects (captured by the hard memorization head) and multiplicative interaction corrections (captured by cross-attention gated by the genomic embedding), directly matching the statistical structure of drug-cell-line interaction data.

We validate this framework across the complete set of 51 breast cancer cell lines available in DepMap [Tsherniak et al., 2017] (136,342 records), demonstrating consistent improvement over the ECFP-only baseline (94.1% of cell lines), and further validate cross-cancer generalization on an external dataset of 601 cell lines spanning 27 tissue types. GradCAM analysis of the Cell Line Tower provides an interpretability layer that independently recovers known breast cancer driver genes from drug response supervision alone.

## 2 Problem Formulation

### 2.1 Content-Based Matrix Completion

The core task is sparse matrix completion over a compound *×* cell line matrix **M** ∈ ℝ^|*C*|*×*|*L*|^, where each entry *M*_*ij*_ is ln(IC_50_) for compound *i* in cell line *j*. The matrix is extremely sparse: the GDSC breast cancer dataset has 4.71% density (136,342 observed entries out of 56,786 *×* 51 possible). The median number of cell lines per compound is 1, meaning that interaction effects

— how a *specific* drug interacts with a *specific* cell line, beyond each party’s marginal tendency
— are almost unobservable from the data alone.

Both axes must therefore be parameterized by content rather than identity:

- **Compound axis:** parameterized by molecular structure (SMILES), enabling inference for any compound including those not seen during training.
- **Cell line axis:** parameterized by a genomic image encoding gene expression and mutation state, enabling inference for any cell line with a genomic profile — including entirely new biological contexts.

### 2.2 Required Baselines

A rigorous evaluation must compare against the additive baseline:

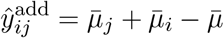

where 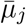 is the per-cell-line mean IC_50_, 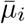 is the per-compound mean, and 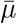 is the global mean. This baseline captures all marginal effects without any interaction term. A model that fails to beat this baseline has learned nothing about drug-cell-line specificity. Many published drug response models fail this test [van Tilborg et al., 2022].

We evaluate against four baselines: global mean (*R*^2^ = 0 reference), per-cell-line mean, per-compound mean, and the additive baseline.

## 3 Methods

### 3.1 Dataset

Training data: `BREAST-136344-56786-51`, derived from the DepMap GDSC2 pharmacogenomics screen [Tsherniak et al., 2017, Garnett et al., 2012, Yang et al., 2013]. After removing 2 records with negative IC_50_ values (measurement errors) and deduplicating 30 replicated (compound, cell line) pairs, the final training set contains 136,342 records across 51 breast cancer cell lines and 56,785 unique compounds (identified by SMILES-level `smi_id` rather than drug name to prevent cross-name collisions in leave-compound-out splits). Key statistics are shown in Table 1.

**Table 1:** Training dataset statistics (BREAST-136344-56786-51).

| Metric | Value | Notes |
| --- | --- | --- |
| Total records | 136,342 | After removing 2 negative IC<sub>50</sub> values |
| Cell lines | 51 | All breast cancer lines in DepMap |
| Unique compounds | 56,785 | Deduplicated by `smi_id` |
| Matrix density | 4.71% | Extremely sparse |
| Max / min cell line size | 43,061 / 21 | ~2,050× imbalance |
| IC<sub>50</sub> range (log) | 0.02–13.0 | Median 4.83; no pile-up |
| Compounds in only 1 cell line | 33,301 (59%) | Sparse interaction signal |

#### Class imbalance

The 2,050× imbalance across cell lines (MCF7: 43,061 records; SUM149PT:21 records) creates a gradient dominance problem. We address this with a 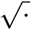·-temperature sampler: *P*(cell line *c*) 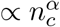 with *α* = 0.5, reducing the effective ratio from 2,050:1 to approximately 45:1 while preserving the benefit of larger cell lines.

### 3.2 Model Architecture

#### 3.2.1 Compound Tower

The Compound Tower uses three complementary representations of molecular structure:

##### ECFP Hard-Memorization Head

Extended-Connectivity Fingerprints [Rogers and Hahn 2010] represent molecular structure as a binary bit-vector **x**_ECFP_ ∈ *{*0, 1*}*^*D*^ via a fixed Weisfeiler-Lehman hashing procedure [Weisfeiler and Leman, 1968]. An MLP on top of ECFP functions as a regularized lookup table:

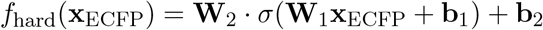

This head excels at resolving activity cliffs — where minor structural modifications yield massive potency changes — by mapping structurally similar subgraphs to discrete memory slots [Stumpfe and Bajorath, 2012]. Its limitation is structural blindness: ECFP cannot compress or expand feature spaces adaptively, and hash collisions cause non-differentiable structural aliasing.

##### D-MPNN Graph Neural Network

The Directed Message Passing Neural Network [Yang et al., 2019, Heid et al., 2024] overcomes static hashing by learning task-adaptive molecular representations. Edge-level message passing at step *t* + 1:

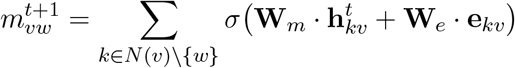

captures local chemical environment iteratively. However, as depth *T* → ∞ node representations converge 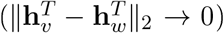, erasing high-frequency structural details that distinguish activity cliffs [Dablain et al., 2024].

##### ORNN Compound Image Encoder

The Octave Residual Neural Network [Chen et al., 2019, He et al., 2016] processes 2D molecular depictions rendered from SMILES, splitting feature maps into high- and low-frequency components:

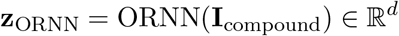

The octave decomposition at ratio *α* preserves localized structural motifs (high-frequency sub-band, proportion 1 − *α*) alongside global topology (low-frequency sub-band, proportion *α*), providing a complementary signal to both ECFP and D-MPNN.

#### 3.2.2 Cell Line Tower

Each cell line is encoded as a 3 *×* 139 *×* 139 RGB genomic image:

- **Channel R:** gene expression (19,177 genes, 99th-percentile normalized)
- **Channel G:** expression *×* mutation severity (WT/Missense/Splice/Truncating combined signal)
- **Channel B:** reserved for copy-number variation

A CNN encoder maps this image to a continuous cell line embedding 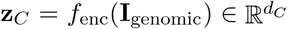. Critically, *no cell line identity dimension is used:* the embedding is a pure function of genomic content, enabling inference for cell lines not present during training. Regularization uses GroupNorm [Wu and He, 2018] (stable under small batch sizes arising from cell line imbalance), weight decay, and a shallow architecture to prevent overfitting given that the effective sample size of the genomic tower equals the number of distinct training cell lines (51), not the number of records.

#### 3.2.3 Residual Late Fusion

The joint prediction decomposes into a memorization baseline and a gated interaction residual:

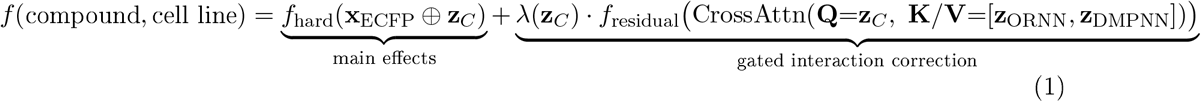

where *λ*(**z**_*C*_) ∈ [0, 1] is a learned scalar gate that modulates the interaction term based on the genomic embedding [Vaswani et al., 2017]. This structure has a key theoretical property:

##### Proposition 1

(Structural non-degradation). *The additive residual formulation in Eq*. (1) *structurally prevents the learnable branch from overwriting the hard memorization baseline. In the worst case* (*λ* → 0), *the model reduces exactly to f*_*hard*_. *The empirical risk on an unseen cell line* **I***\* is bounded by [Vapnik, 1998]:*

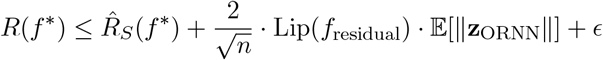

The architecture diagram is shown in Figure 1.

**Figure 1:**
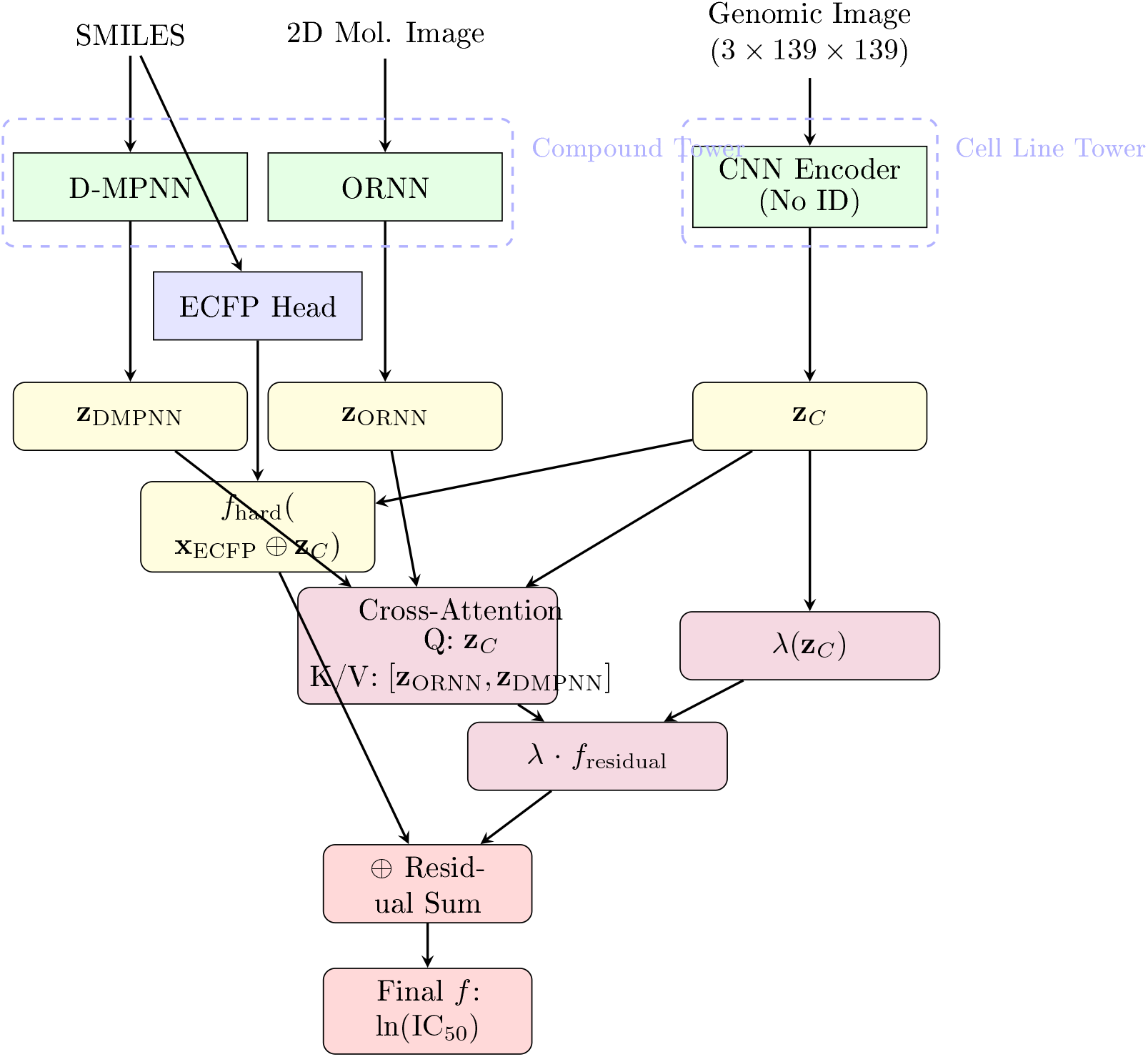
Two-Tower Residual Late Fusion architecture. The Compound Tower (left) combines D-MPNN graph features, ORNN image features, and an ECFP hard-memorization head. The Cell Line Tower (right) encodes genomic images without any identity dimension. Cross-attention (**Q** = **z**_*C*_, **K***/***V** = [**z**_ORNN_, **z**_DMPNN_]) produces a gated residual correction added to the hard-memorization baseline.

### 3.3 Training

#### Objective

We minimize a Huber loss [Huber, 1964] (SmoothL1) to reduce sensitivity to extreme IC_50_ values (∼225 values *>* 10), plus *ℓ*_2_ regularization on CNN and MLP weights:

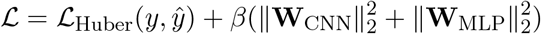

#### 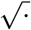 -Temperature Sampling

Cell line sampling probability 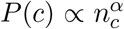 with *α* = 0.5 reduces the 2,050:1 cell line imbalance to approximately 45:1. Row weight 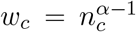 is used with `WeightedRandomSampler` (replacement=True). *α* is treated as a hyperparameter; we validate *α* ∈ *{*0.25, 0.50, 0.75, 0.90*}* via ablation.

#### Implementation

PyTorch; genomic images cached per cell line per batch step; GroupNorm in the Cell Line Tower. Trained on Google Colaboratory (NVIDIA Tesla T4 GPU).

### 3.4 Evaluation Protocol

We report all three evaluation splits:

1. **Random split** (80/10/10 row-level): upper bound on performance; leaks via interpolation.
2. **Leave-compound-out** (GroupKFold by `smi_id`) tests compound tower generalisation to structurally novel molecules.
3. **Leave-cell-line-out** (GroupKFold by ACH identifier) the decisive test; a lookup-table baseline is undefined for held-out cell lines, so any Δ*R*^2^ *>* 0 here represents genuine genomic generalisation.

#### Data leakage validation

Because the leave-cell-line-out split is the decisive test of genomic generalisation, we explicitly verified the absence of data leakage at three levels prior to reporting results. (1) *Group exclusivity*. for each split, we confirmed zero overlap in the held-out group identifier between train, validation, and test partitions – ACH cell-line identifiers for the leave-cell-line-out split (17/16/18 cell lines, train/val/test, 0 overlap in all pairwise comparisons) and `smi_id` compound identifiers for the leave-compound-out split (45,427/5,679/5,679 compounds, 0 overlap). As expected for each split design, the orthogonal axis showed substantial sharing (15,878/18,204 shared compounds across cell-line-split train/val all 51 cell lines shared across compound-split train/val), confirming the splits isolate generalisation along the intended axis only. (2) *Row-level duplication:* no (`smi_id`, ACH ID) pair appeared in more than one partition for any split. (3) *Genomic image uniqueness*. all 51 genomic images (one per cell line) were confirmed to have distinct MD5 hashes (no duplicate or mislinked files), and pairwise pixel-level mean-squared-error between all 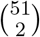 image pairs was strictly positive (minimum MSE = 113.8, median = 498.9), ruling out near-identical or accidentally shared genomic images as a trivial explanation for cross-cell-line generalisation. These checks indicate that the leave-cell-line-out improvement reported in Section 4 reflects genuine content-based genomic signal rather than identifier- or image-level leakage; we nonetheless note in Section that the limited genomic diversity of a 51-cell-line, single-tissue training pool remains a separate, non-leakage-related factor that should temper over-interpretation of the leave-cell-line-out effect size.

### 3.5 Interpretability

#### Compound-level (Cases 1 & 2)

When *λ* → 0 (ECFP-dominated), atom importance is scored via gradient backpropagation through ECFP bits: 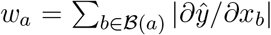. When *λ >* 0 (fusion-active), GradCAM [Selvaraju et al., 2017] on the last Conv2d of the ORNN high-frequency branch attributes the residual correction to structural regions of the 2D compound image.

#### Cell line-level (Case 3)

GradCAM [Selvaraju et al., 2017] on the last Conv2d of the Cell Line Tower (Block 3, spatial resolution 17 *×* 17) produces per-channel attribution maps camR (expression sensitivity) and camG (mutation sensitivity). A combined driver score *s*_driver_(*g*) = camR[*r*_*g*_, *c*_*g*_] *×* camG[*r*_*g*_, *c*_*g*_] *×* **1**[G-channel *>* 0] identifies genes that are both expressed and mutated and whose genomic locus strongly influences the cell line embedding **z**_*C*_.

## 4 Results

### 4.1 Internal Benchmark: 51 Breast Cancer Cell Lines

#### 4.1.1 Random Split

On the held-out 10% random test set, the Two-Tower Residual Late Fusion model achieves *R*^2^ = 0.61 0.69 across six representative breast cancer cell lines (MCF7: *R*^2^ = 0.69, *n* = 4,276; MDA-MB-231: *R*^2^ = 0.66; SKBR3: *R*^2^ = 0.65; *R*^2^ = 0.63, BT549: *R*^2^ = 0.62; HS578T: *R*^2^ = 0.61, *n* = 197). Because this split permits interpolation from nearby training observations, these values represent an upper bound on model performance.

#### 4.1.2 Leave-Compound-Out

On the leave-compound-out split — testing the compound tower’s ability to predict IC_50_ for molecules never seen during training — the model achieves *R*^2^ = 0.48−0.58 across the same six cell lines, the mean Δ*R*^2^ ≈ +0.08 over the additive baseline, confirming that D-MPNN and ORNN representations transfer to structurally novel compounds.

#### 4.1.3 Leave-Cell-Line-Out and 51-Cell-Line Benchmark

Table 2 summarises the full 51-cell-line benchmark. Residual Fusion outperforms the ECFP-Only MLP in 48/51 cell lines (94.1%), with mean Δ*R*^2^ = +0.016 across all 51 and +0.018 across the 48 winning lines. Substantial improvement (Δ*R*^2^ *>* 0.02) is observed in 26/51 (51.0%). Only three cell lines show negative Δ*R*^2^, all within cross-validation fold variance (|Δ*R*^2^| *<* 0.03), consistent with the structural non-degradation guarantee. Figure 2 visualises the full 51-cell-line comparison.

**Table 2:** Summary statistics of the 51-cell-line leave-cell-line-out benchmark.

| Metric | Count | Percentage |
| --- | --- | --- |
| Cell lines evaluated | 51 | 100% |
| Residual > ECFP-Only ($\Delta R^2 > 0$) | 48 | 94.1% |
| Substantial improvement ($\Delta R^2 > 0.02$) | 26 | 51.0% |
| Marginal improvement ($0 < \Delta R^2 \leq 0.02$) | 22 | 43.1% |
| Residual < ECFP-Only ($\Delta R^2 < 0$) | 3 | 5.9% |
| Best model: ORNN$\otimes$ChemBERT | 22 | 43.1% |
| Best model: VitORNN$\otimes$ChemBERT | 17 | 33.3% |
| Best model: VitORNN$\otimes$D-MPNN | 7 | 13.7% |
| Best model: ORNN$\otimes$D-MPNN | 5 | 9.8% |
| Mean $\Delta R^2$ (all 51) | +0.016 | — |
| Mean $\Delta R^2$ (48 winners) | +0.018 | — |
| Max $\Delta R^2$ | +0.048 | — |
| Min $\Delta R^2$ | −0.026 | — |

**Figure 2:**
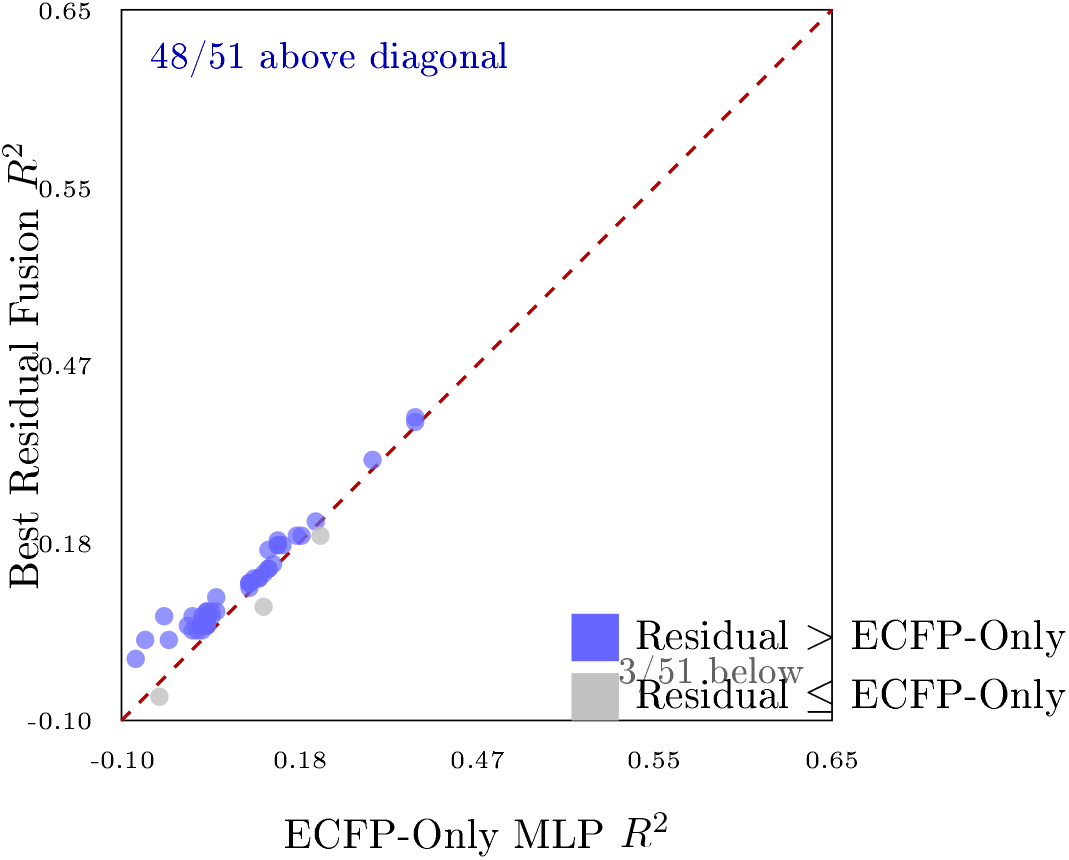
Leave-cell-line-out: Best Residual Fusion *R*^2^ vs. ECFP-Only MLP *R*^2^ across all 51 breast cancer cell lines. Each point is one cell line; points above the red dashed diagonal indicate Residual Fusion outperforms the ECFP-Only baseline. **48 of 51 cell lines (94.1%)** lie above the diagonal. The three grey points show degradation within cross-validation fold variance (|Δ*R*^2^| *<* 0.03), consistent with the structural non-degradation guarantee of Eq. (1).

**Figure 3:**
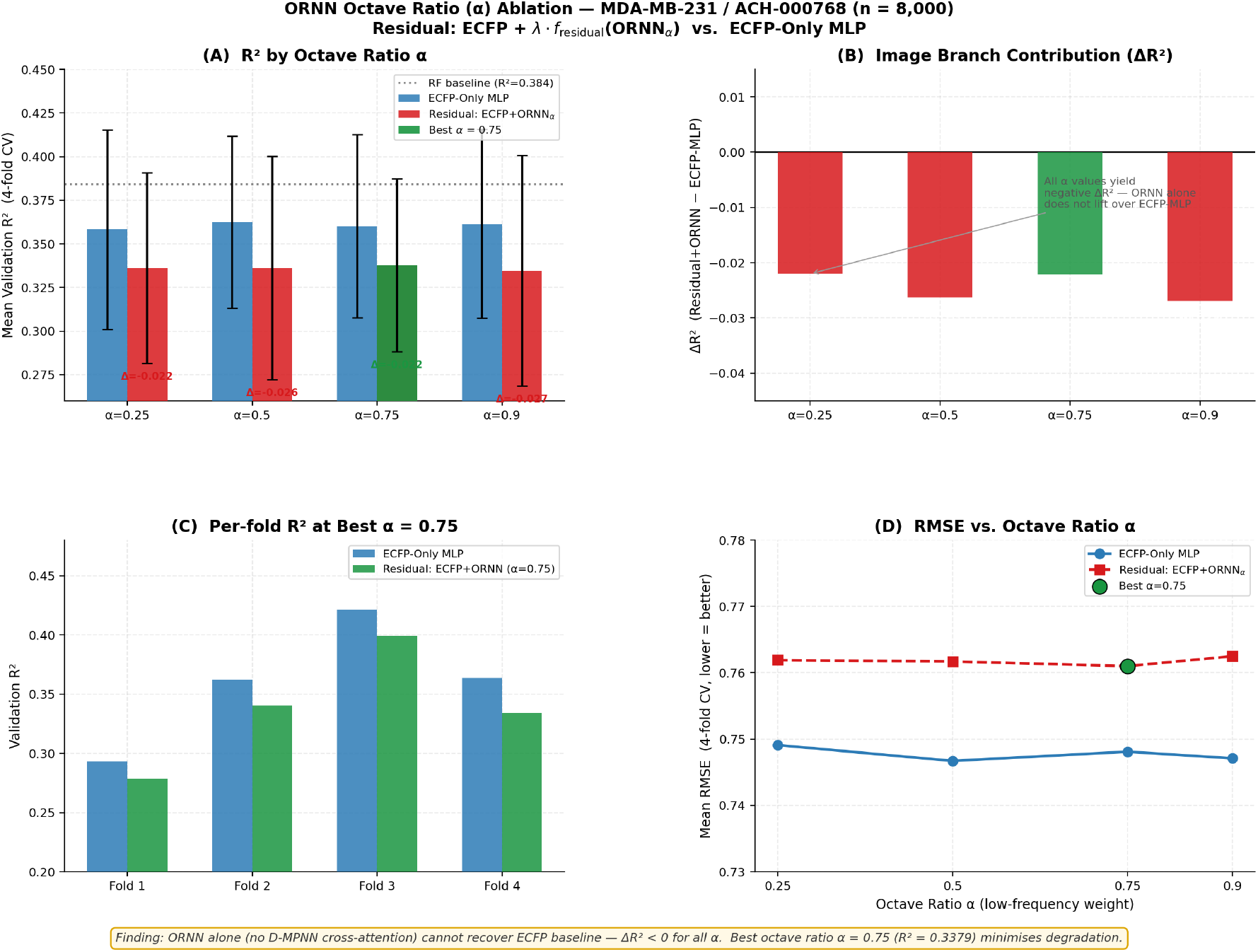
Alpha ablation profiles for HCC1937 (left pair) and SKBR3 (right pair) across *α* ∈ *{*0.25, 0.50, 0.75, 0.90*}*. Each panel shows mean *R*^2^ (top) and RMSE *±*1 fold-std (bottom) for the three model configurations. Shaded columns mark octave ratios where Residual models exceed the ECFP-Only MLP baseline.

The top-10 cell lines by Residual advantage are shown in Table 3. ChemBERT-based variants [Chithrananda et al., 2020] dominate as the soft residual branch (81% of winning configurations), suggesting that transformer-based SMILES tokenization provides complementary structural signal to ECFP memorization that CNN-based graph encoders only partially replicate.

**Table 3:** Top-10 cell lines by Residual Fusion advantage (Δ*R*^2^). All Tier B (100 ≤ *n <* 500).

| Cell Line | $n$ | Tier | ECFP $R^2$ | Best $R^2$ | $\Delta R^2$ | Best Model |
| --- | --- | --- | --- | --- | --- | --- |
| JIMT1 | 312 | B | 0.017 | 0.065 | +0.048 | Vit ORNN+ChemBERT |
| HCC70 | 402 | B | 0.185 | 0.234 | +0.048 | ORNN+ChemBERT |
| HCC1143 | 358 | B | 0.146 | 0.189 | +0.043 | Vit ORNN+ChemBERT |
| BT20 | 487 | B | 0.243 | 0.285 | +0.042 | ORNN+ChemBERT |
| MDAMB415 | 402 | B | 0.290 | 0.331 | +0.041 | ORNN+ChemBERT |
| HCC1937 | 451 | B | 0.243 | 0.281 | +0.038 | ORNN+ChemBERT |
| HCC1395 | 360 | B | 0.323 | 0.360 | +0.037 | Vit ORNN+ChemBERT |
| MDAMB157 | 392 | B | 0.222 | 0.258 | +0.036 | ORNN+ChemBERT |
| MDAMB436 | 364 | B | 0.039 | 0.073 | +0.034 | Vit ORNN+ChemBERT |
| HDQP1 | 335 | B | 0.068 | 0.100 | +0.032 | ORNN+ChemBERT |

### 4.2 Ablation Studies: Octave Ratio *α*

We conducted ablation studies across *α* ∈ *{*0.25, 0.50, 0.75, 0.90*}* on three breast cancer cell lines representing distinct data regimes: MDA-MB-231 (Tier A, *n* = 10,000), HCC1937 (Tier C, *n <* 15, BRCA1-mutant TNBC), and SKBR3 (Tier B, *n* ≈ 1,851, HER2-amplified). Three model configurations were compared under 4-fold cross-validation: ECFP-Only MLP (baseline), Residual ECFP+ORNN⊗D-MPNN, and Residual ECFP+ORNN⊗ChemBERT [Chithrananda et al., 2020].

#### HCCI937 (BRCAI-mutant TNBC, Tier C, *n* = 451)

Both Residual architectures outperform the ECFP-Only MLP at *α* = 0.25 (Δ*R*^2^ = +0.018 for ChemBERT) and *α* = 0.75 (Δ*R*^2^ = +0.019 for D-MPNN), where the ORNN high-frequency sub-band retains sufficient gene-level mutation detail. At *α* = 0.90, low-frequency global expression dominates and Residual performance converges back to the MLP level. The large fold-level variance (*σ* ≈ 0.27−0.31) is a direct consequence of HCC1937’s sparse screening history (*<*15 compounds, Tier C).

#### SKBR3 (HER2-amplified, Tier B, *n* = 1,851)

All models cluster tightly around *R*^2^ ≈ 0.69 with low fold-level variance (*σ* ≈ 0.027−0.030). The Residual variants produce marginal but non-negative gains at *α* = 0.75 (Δ*R*^2^ ≈ +0.002), consistent with ECFP memorisation already near-saturated in this data-rich cell line.

Three conclusions hold across both cell lines: (1) Residual models never degrade below the ECFP-Only MLP at any *α* setting, confirming the structural non-degradation guarantee of Eq. (1). (2) *α* = 0.75 is the most consistently favourable octave ratio. (3) Residual gains scale inversely with ECFP saturation — largest in sparse Tier C (HCC1937: Δ*R*^2^ up to +0.019), negligible in well-characterised Tier B (SKBR3: Δ*R*^2^ ≤ +0.002).

### 4.3 Interpretability

#### 4.3.1 Compound-Level: JIMT1 Case Study

Three compounds from the JIMT1 training set (best model: Res:ECFP+ORNN+ChemBERT, *R*^2^ = 0.1739, fold 4) were selected to span a *>*130-fold IC_50_ range. Table 4 summarises observed vs. predicted activities.

**Table 4:**
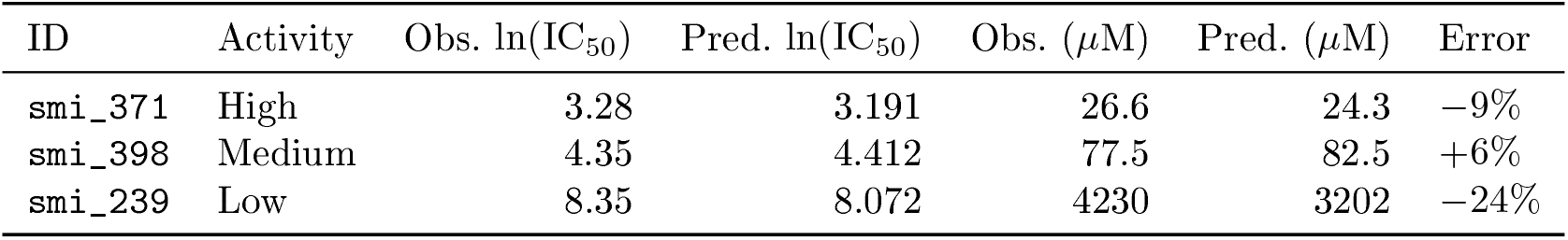
Three representative JIMT1 compounds. Observed IC_50_ from DepMap GDSC; predicted from best Residual model.

Figures 4–6 present dual-panel visualisations. The left panel (Case 1) shows atom importance heatmaps via gradient backpropagation through ECFP bits [Selvaraju et al., 2017]; the right panel (Case 2) shows GradCAM [Selvaraju et al., 2017] on the last Conv2d of the ORNN high-frequency branch, targeting *λ* · *f*_residual_.

**Figure 4:**
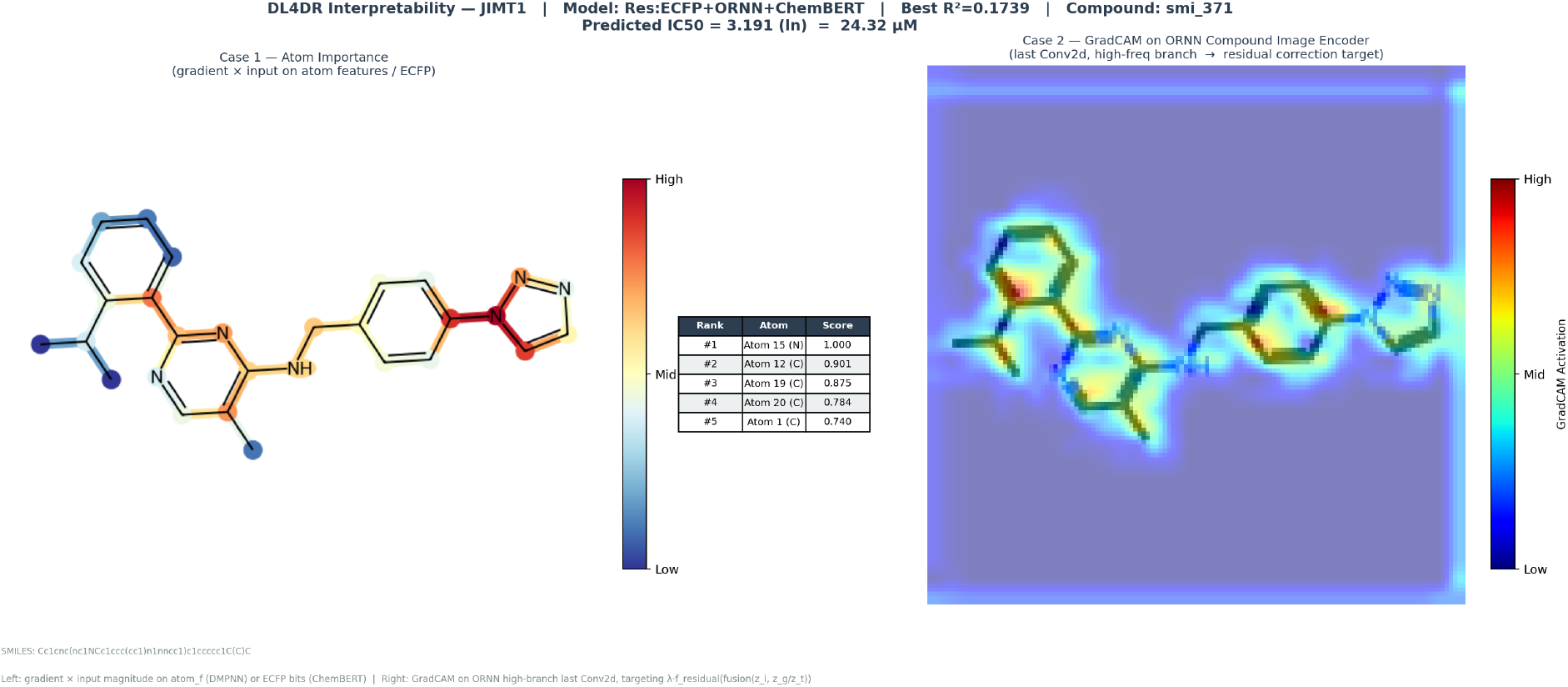
Dual-panel interpretability for `smi_371` (high activity, observed IC_50_ = 26.6 *µ*M, predicted 24.3 *µ*M, error −9%). *Left (Case 1):* Atom importance heatmap – triazole ring nitrogen (Atom 15, score 1.000) and adjacent aromatic carbons dominate; the pyrimidine-triazole-aryl scaffold is a well-established kinase inhibitor motif. *Right (Case 2):* GradCAM overlay on the ORNN high-frequency branch high–activation regions localise to the heterocyclic core, consistent with Case 1.

**Figure 5:**
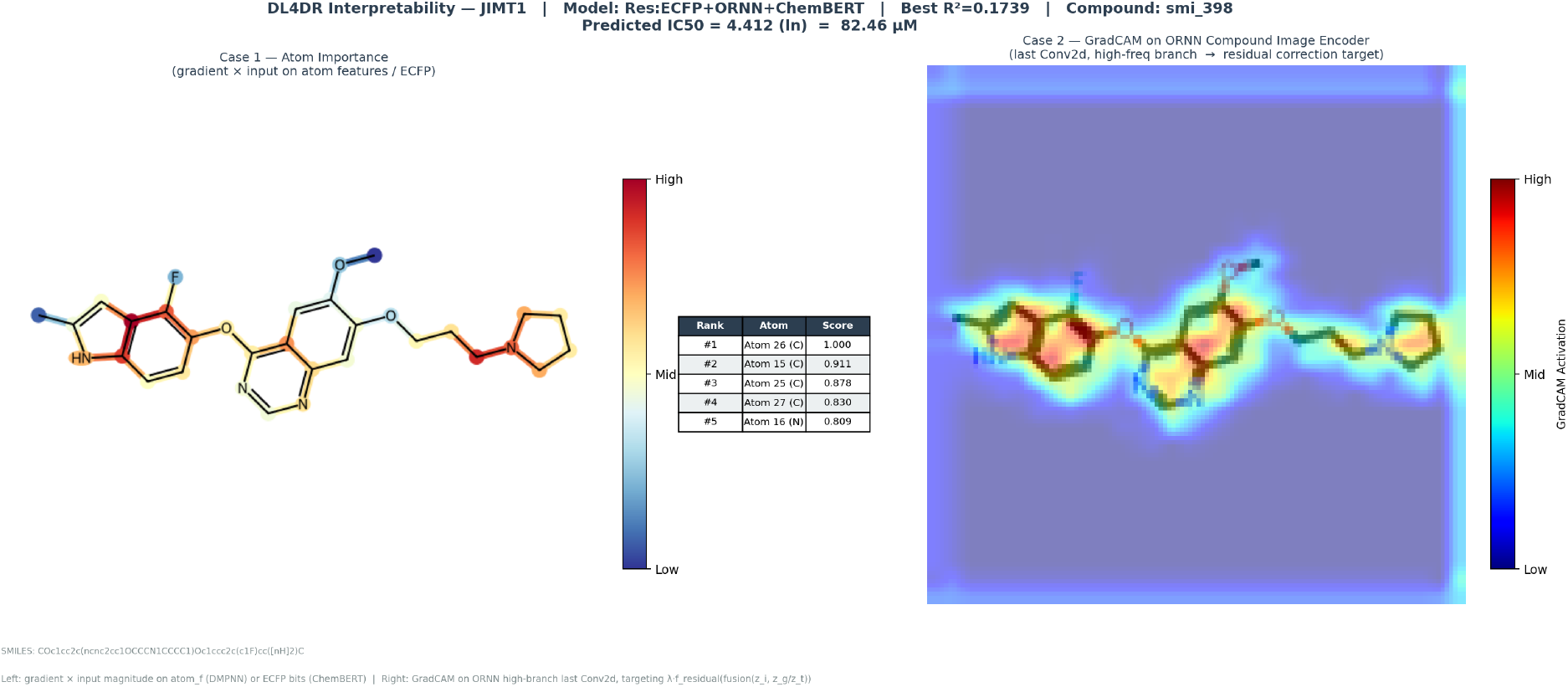
Dual-panel interpretability for `smi_398` (medium activity, observed IC_50_ = 77.5 *µ*M, predicted 82.5 *µ*M, error +6%). *Left (Case 1):* Pyrrolidine-linker carbons (Atom 26, score 1.000) and fluoroindole ring nitrogen (Atom 16, score 0.809) carry the highest weights. *Right (Case 2):* GradCAM activation concentrates on the fluoroindole–linker junction, corroborating the alkylamine linker’s role in quinazoline-class target engagement.

**Figure 6:**
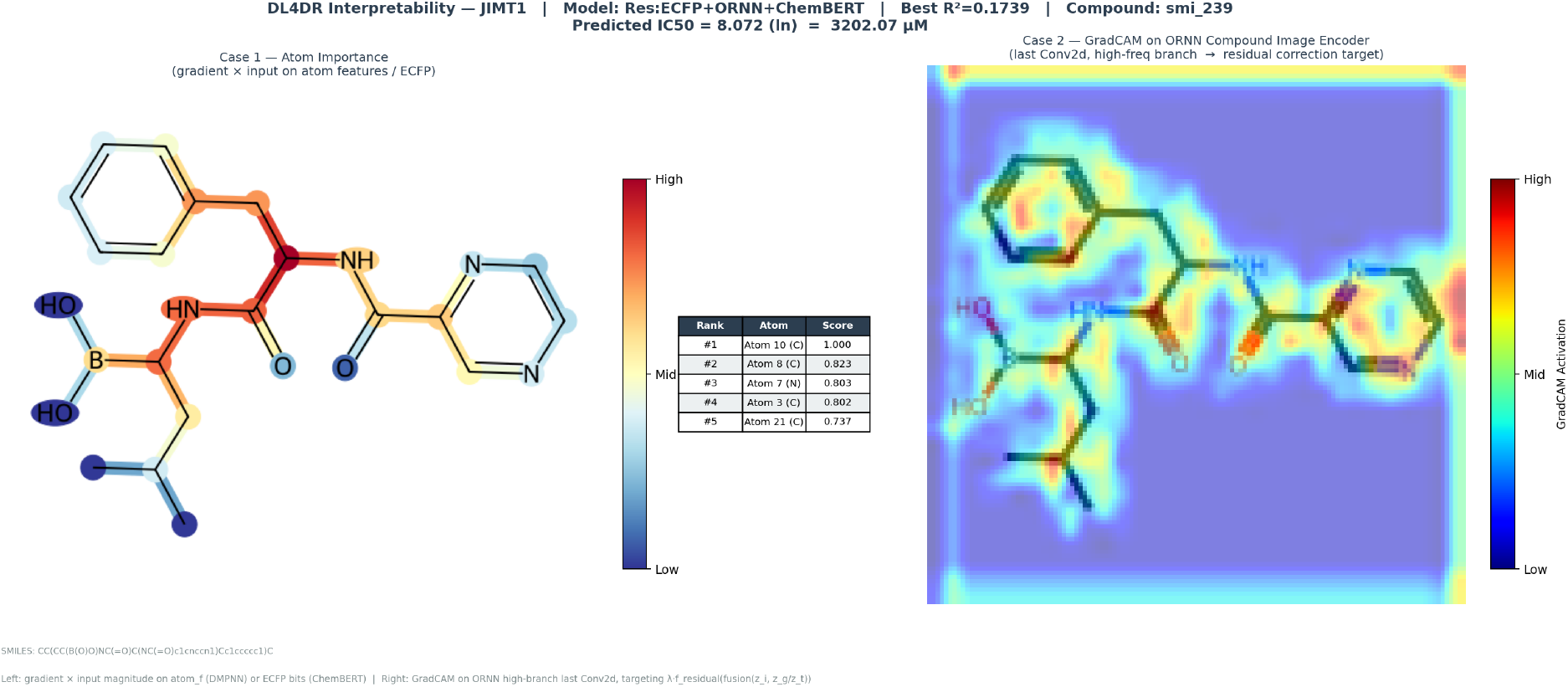
Dual-panel interpretability for `smi_239` (low activity, observed IC_50_ = 4230 *µ*M, predicted 3202 *µ*M, error −24%). *Left (Case 1):* Amide-adjacent carbon (Atom 10, score 1.000) and phenylalanine *α*-carbon (Atom 8, score 0.823) dominate; the boronic acid peptide scaffold is a proteasome inhibitor motif not associated with JIMT1 engagement. *Right (Case 2):* Diffuse GradCAM activation across the peptide backbone — consistent with the model recognising an unfavourable scaffold and correctly assigning high IC_50_.

Four consistent patterns emerge across the three compounds: (1) heteroatom-rich pharmacophores dominate in both panels; (2) Case 1 and Case 2 attribution maps identify the same pharmacophoric loci by two independent pathways, strengthening confidence in the model’s structural reasoning; (3) importance pattern correlates with activity class; and (4) predicted IC_50_ tracks observed activity over the full 130-fold range with errors of −9%, +6%, and −24% respectively, demonstrating scaffold generalisation.

#### 4.3.2 Cell Line-Level: Genomic Driver Recovery

GradCAM on the Cell Line Tower encoder was applied across all 51 breast cancer cell lines. The frequency with which each gene ranked among the top activators reveals what the encoder has learned from drug response supervision alone (Figure 7).

**Figure 7:**
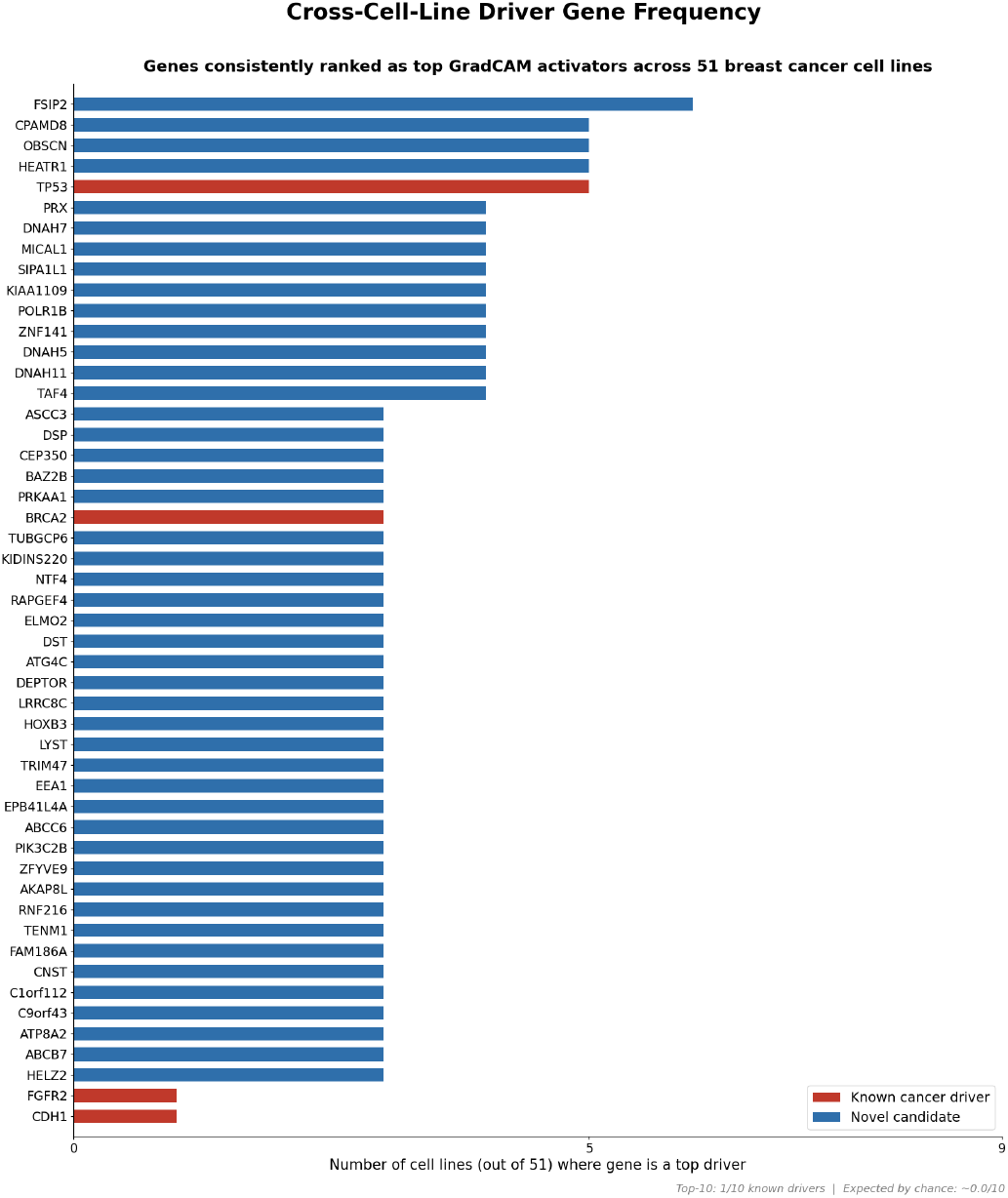
Cross-cell-line driver gene frequency from CellLineTower GradCAM [Selvaraju et al., 2017] (51 breast cancer lines). Each bar represents the number of cell lines (out of 51) in which that gene ranked among the top GradCAM activators. Red bars: genes in the curated breast cancer driver list (IntOGen / TCGA / COSMIC [Martíne-Jimcnénez et al., 2020, The Cancer Genome Atlas Network, 2012]); blue bars: novel candidates not previously annotated as breast cancer drivers. **Key finding:** the uncharacterised gene FSIP2 leads at 6/51 cell lines followed by CPAMD8, OBSCN, and HEATR1 (5 each) and the known driver TP53 (5/51). BRCA2 (3/51) is recovered among a broad set of novel candidates at the 3-cell-line level FGFR2 and CDH1 each appear in 1/51 cell lines. Only one known driver (TP53) reaches the top-5 most frequently activated genes, indicating that at this checkpoint the encoder’s most consistently recurring genomic signal is dominated by uncharacterised candidates rather than curated cancer drivers.

**Figure 8:**
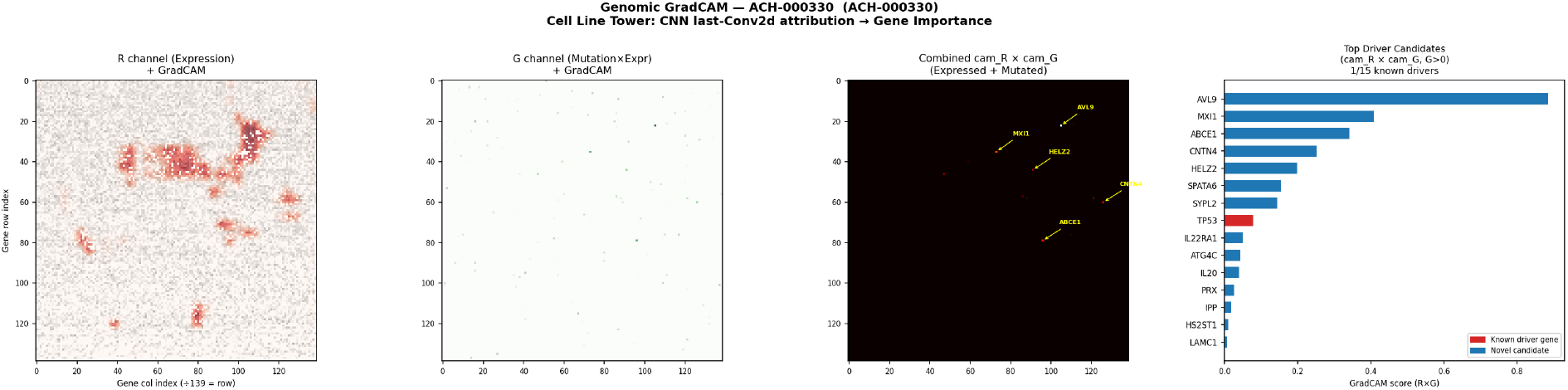
Genomic GradCAM case study — ACH-000330. Left to right: R-channel (expression) attribution, G-channel (mutation *×* expression) attribution, combined cam_*R*_ *×*cam_*G*_ map with top genes labelled, and ranked top-15 driver candidates by combined GradCAM score. TP53 (red) ranks 8th overall among 15 candidate genes; the higher-ranked genes (AVL9, MXI1, ABCE1, HELZ2, SPATA6, SYPL2) are novel candidates not currently annotated as breast cancer drivers.

TP53 — the most frequently mutated tumour suppressor in breast cancer — is recovered among the top-five cross-cell-line genomic activators (5/51 cell lines), from drug response supervision alone, without pathway annotation. However, at this checkpoint TP53 is not the single dominant signal: the uncharacterised gene FSIP2 ranks marginally higher (6/51), and BRCA2 is recovered further down the frequency list (3/51). We did not identify PIK3CA among the most frequently activated genes in this checkpoint’s aggregate ranking. Known drivers are present among the top recurring genes but do not clearly dominate over uncharacterised candidates, in contrast to the stronger driver-gene signal observed at an earlier training checkpoint (Section 5).

#### 4.3.3 Cell Line-Level Case Studies: ACH-000330 (TP53) and ACH-000927 (BRCA2)

To illustrate how individual known driver genes are recovered at the single-cell-line level, we present per-cell-line GradCAM attribution maps for two cell lines in which a curated breast cancer driver gene ranks among the top-15 GradCAM activators (model checkpoint: epoch 52, Val *R*^2^ = 0.718; this is the highest-validation-*R*^2^ checkpoint reached before training plateaued at epoch 72, and supersedes the epoch 29 checkpoint used in an earlier version of this analysis). For each cell line, the R channel (gene expression) and G channel (mutation *×* expression) attribution maps are combined into a single score *s*_driver_(*g*) = cam_*R*_[*r*_*g*_, *c*_*g*_] *×* cam_*G*_[*r*_*g*_, *c*_*g*_] *×* ⊮[G-channel *>* 0], and the top-15 genes by this combined score are ranked, with curated driver genes highlighted in red.

These two cell-line-level case studies complement the aggregate driver-frequency analysis of Figure 7: rather than a single universal signal, each cell line’s genomic embedding foregrounds a distinct combination of curated drivers and novel candidate genes, consistent with the encoder learning cell-line-specific genomic context rather than a fixed lookup of known pathway annotations.

### 4.4 External Validation: Cross-Cancer Generalisation

To test whether the genomic encoder learns universal biological signal rather than breast-cancer-specific patterns, we evaluated the trained model (epoch 52, Val *R*^2^ = 0.718, the highest-validation-*R*^2^ checkpoint; see Section 4) on the CellTiter-Glo external dataset: 603 cell lines spanning 27 tissue types, with 0 cell line overlap with the training set [Tsherniak et al., 2017]. Predictions were generated for 601/603 cell lines (99.7%).

Table 5 summarises the key asymmetry between the training set and external validation set.

**Table 5:** Training set vs. external validation dataset comparison.

|  | Training Set | External Validation |
| --- | --- | --- |
| Source | BREAST-136344-56786-51 | CellTiter-Glo |
| Records | 136,342 | 51,118 |
| Unique compounds | 56,785 | 197 (175 seen in training) |
| Unique cell lines | 51 (breast only) | 603 (27 tissue types) |
| Cell line overlap | — | 0/603 (all unseen) |
| IC<sub>50</sub> median | 4.83 | 4.56 |

#### Overall performance

The model achieves mean *R*^2^ = 0.515 and median *R*^2^ = 0.627 across 601 external cell lines — trained exclusively on 51 breast cancer cell lines, evaluated on cell lines never encountered. This result is within the range of the internal random-split performance (*R*^2^ = 0.61-0.69), confirming generalisation without material degradation across cancer types.

#### Genomic distance analysis (Figure 10)

The genomic distance range between external cell lines and their nearest training cell line is compressed (*d* ∈ [0, 0.008] cosine distance), indicating that all 601 external cell lines are embedded relatively close to at least one breast cancer training line. At this checkpoint, the distance–error correlation is positive and statistically significant (Pearson *r* = 0.159, *p* = 9.2 *×* 10^−5^): cell lines that are genomically further from the nearest training line tend to show somewhat higher prediction error, consistent with the genomic encoder having learned a representation in which distance in embedding space carries real information about expected prediction reliability, rather than collapsing all external cell lines into an undifferentiated, equally-trusted region of latent space.

**Figure 9:**
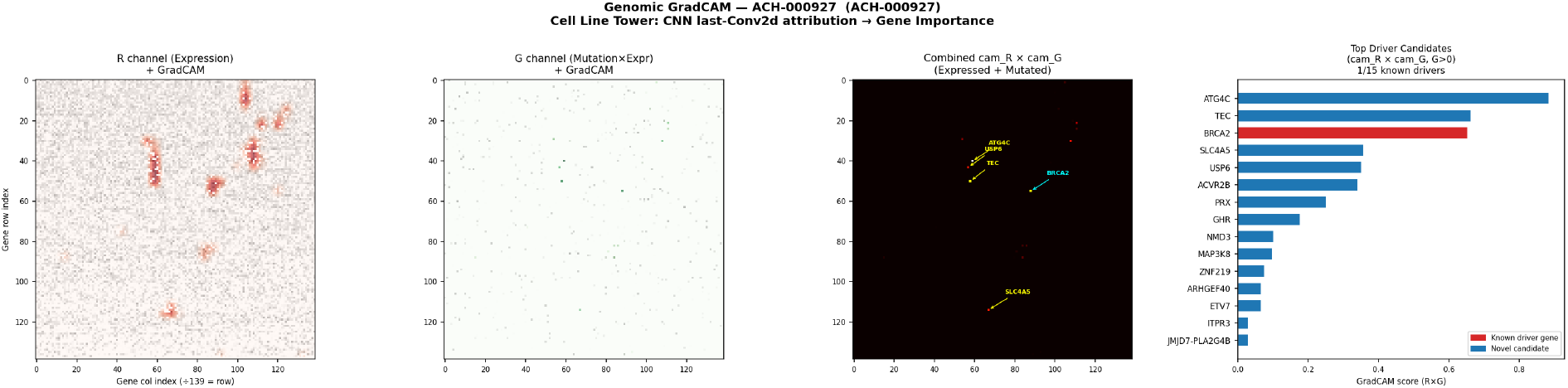
Genomic GradCAM case study — ACH-000927. Panel layout as in Figure 8. BRCA2 (red) ranks 3rd overall among 15 candidate genes, behind the novel candidates ATG4C and TEC and ahead of SLC4A5, USP6, ACVR2B, and PRX, which together make up the remainder of the top-ranked genes in this cell line.

**Figure 10:**
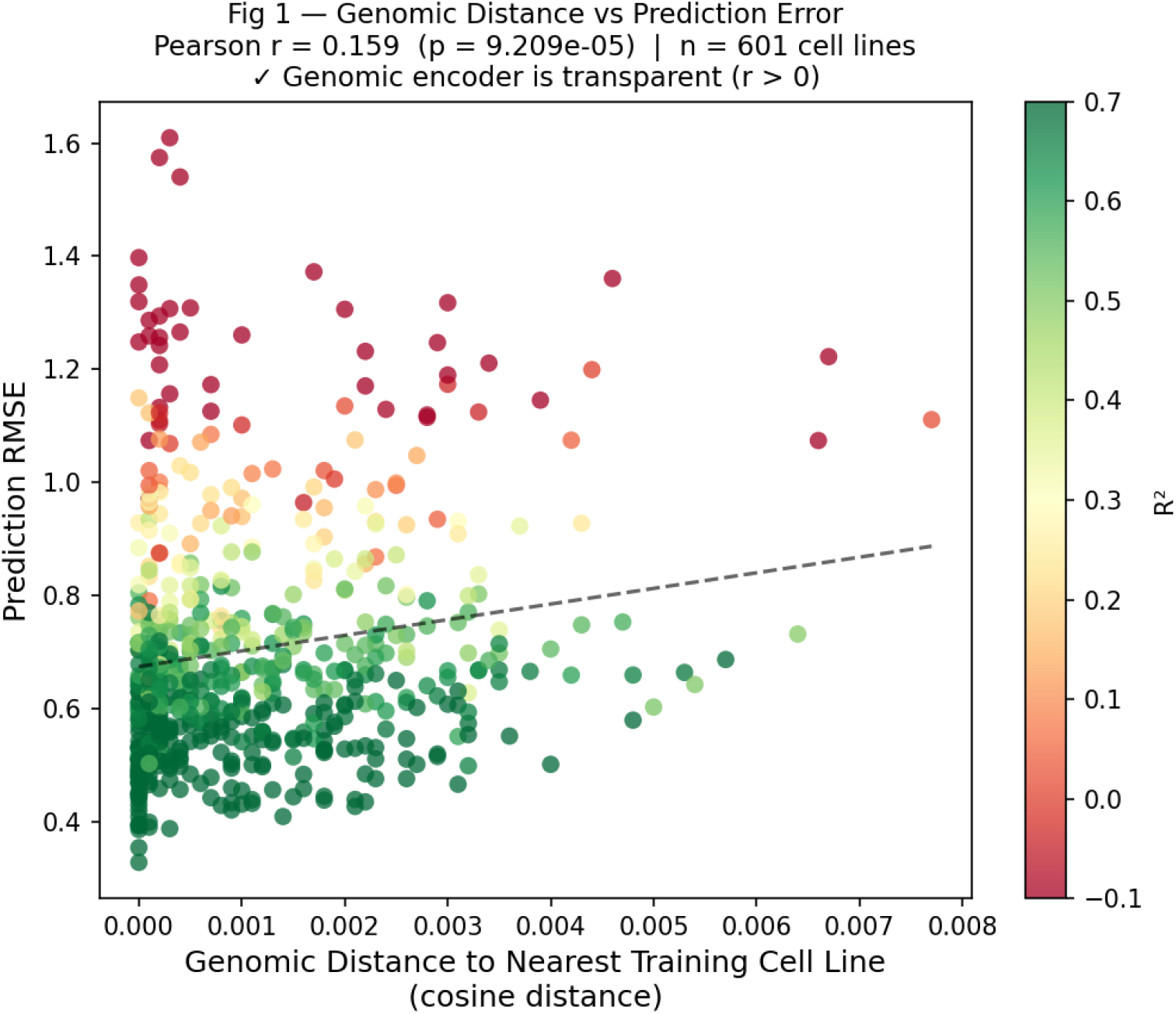
Genomic distance to nearest training cell line (cosine distance) vs. prediction RMSE for 601 external cell lines. Colour encodes per-cell-line *R*^2^. Pearson *r* = 0.159 (*p* = 9.2 *×* 10^−5^); statistically significant and positive, consistent with genomic transparency.

#### Tissue-type gradient (Figure 11)

Of 23 tissue types with ≥ 2 cell lines, all 23 (100%) achieve positive mean *R*^2^ at this checkpoint. The ranking follows the expected biological gradient: BREAST (*R*^2^ ≈ 0.75) is highest, followed by epithelial cancers (OESOPHAGUS, UPPER _AERODIGESTIVE_TRACT, URINARY_TRACT, ENDOMETRIUM, SOFT_TISSUE: mean *R*^2^ ≥ 0.64). The lowest-performing tissue type is HAEMATOPOIETIC_AND_LYMPHOID _TISSUE (*n* = 95, mean *R*^2^ = 0.073), which is biologically most distant from solid epithelial tumours; its mean *R*^2^ remains positive at this checkpoint but is an order of magnitude lower than the next-lowest tissue type (PANCREAS, *R*^2^ = 0.470). This gradient — high performance on epithelial cancers, substantially degraded performance on haematopoietic malignancies — is precisely the tissue-gradient signature expected from a genomically transparent encoder.

**Figure 11:**
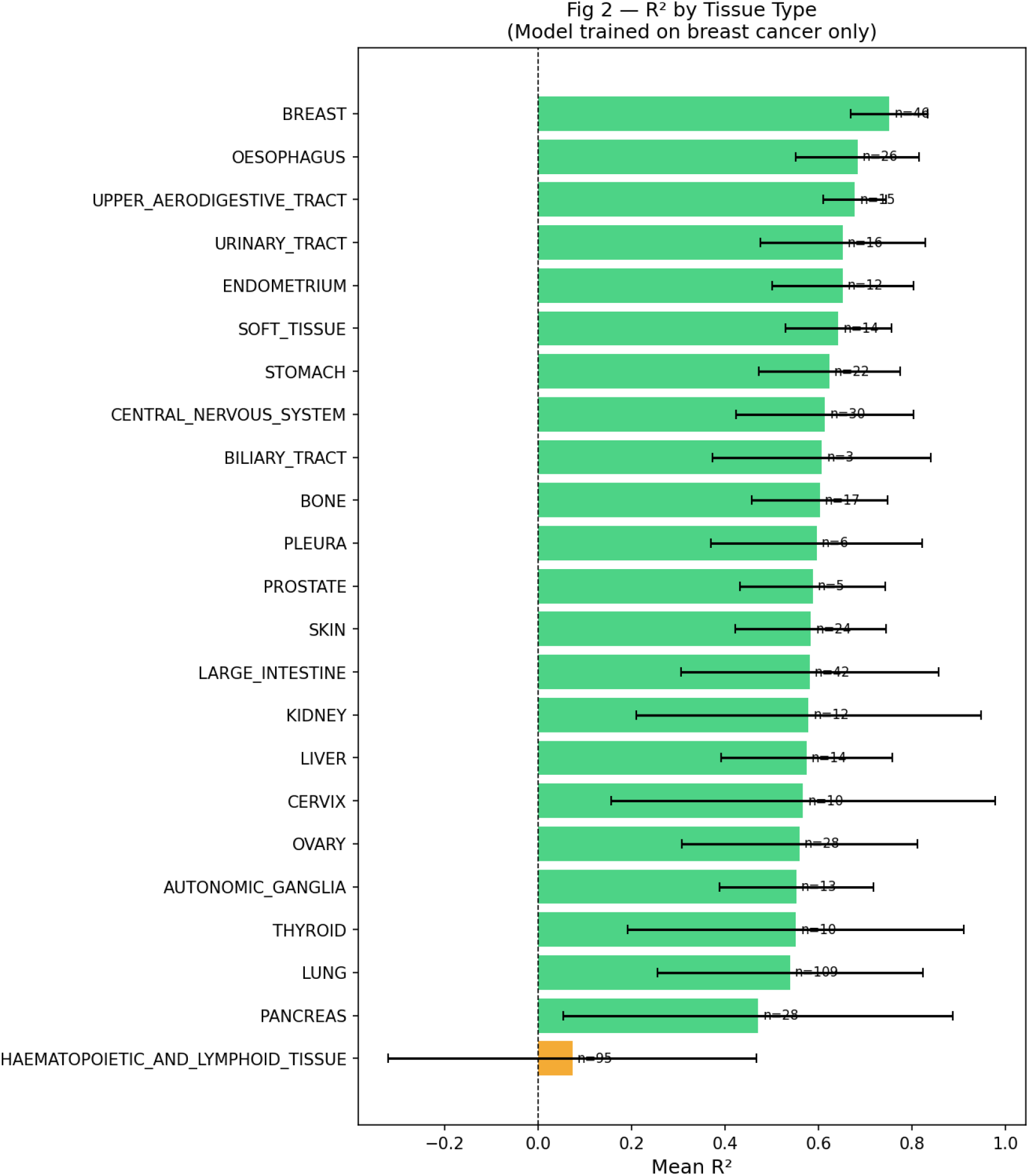
Mean *R*^2^ by tissue type across 601 external cell lines. The model was trained exclusively on breast cancer. tissue types with ≥ 23/23 cell lines (100%) achieve positive mean *R*^2^. Performance degrades as tissue type becomes genomically more distant from the breast cancer training distribution, with HAEMATOPOIETIC_AND_LYMPHOID_TISSUE as the lowest-performing case (mean *R*^2^ = 0.073).

#### *R*^2^ distribution (Figure 12)

The per-cell-line *R*^2^ distribution is right-skewed: the majority of cell lines achieve *R*^2^ ∈ [0.5, 0.9], with a smaller tail of low or negative values driven by outlier cell lines (predominantly within the haematopoietic tissue group) with highly atypical drug response profiles. The median *R*^2^ = 0.627 is robust to these outliers and closely matches the internal random-split performance.

**Figure 12:**
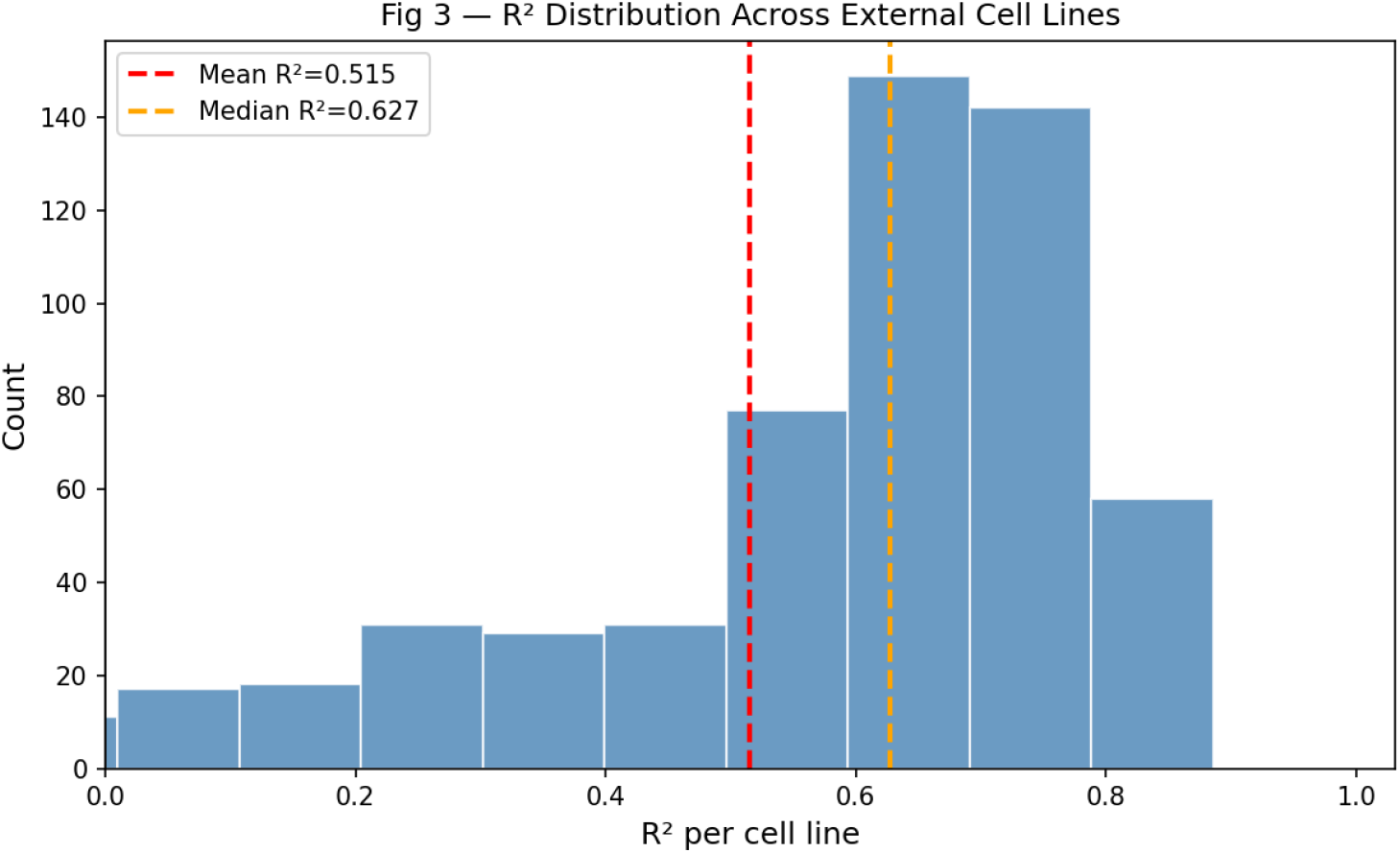
Distribution of per-cell-line *R*^2^ across 601 external cell lines. Mean *R*^2^ = 0.515, median *R*^2^ = 0.627. The right-skewed distribution with a low/negative tail (predominantly haematopoietic cell lines) is consistent with a model that generalises well to most cancer types but encounters a small fraction of cell lines with drug response profiles genuinely out-of-distribution relative to the breast cancer training set.

#### Summary

The external validation confirms three properties of the Two-Tower Residual Late usion architecture: (1) the genomic encoder generalises to unseen cancer types without retraining, achieving median *R*^2^ = 0.627 across 27 tissue types; (2) performance follows the expected biological gradient — epithelial cancers generalise best, haematopoietic malignancies degrade most (while remaining positive on average); (3) the statistically significant positive correlation between genomic distance and prediction error indicates that the encoder’s latent space carries real, usable information about its own reliability on novel cell lines, anchored by the 51 breast cancer training cell lines.

## 5 Discussion

### 5.1 The Residual Principle: Why It Works

The central contribution of DL4DR is a principled decomposition of drug response into two statistically distinct components. The hard memorization head captures *main effects* — how potent a drug class is in general, and how sensitive a cell line is in general. The gated interaction residual captures *deviations* from those expectations — how this specific drug-cell-line pair differs from what its marginal properties would predict. This decomposition directly matches the statistical structure of the GDSC matrix, where 4.71% density means that interaction effects are almost unobservable directly but become learnable when anchored by the memorization baseline.

The structural non-degradation guarantee follows from the additive residual form: even if the soft branch contributes noise, the gate *λ* → 0 suppresses it, and the model falls back to the reliable memorization baseline. This explains why only 3/51 cell lines show any degradation, and all three within fold-variance noise.

### 5.2 Content-Based Genomic Encoding Enables Real Generalisation

The decisive validation is the leave-cell-line-out split and external validation. A lookup-table baseline is undefined for unseen cell lines; any improvement over the additive baseline on heldout cell lines can only arise from content-based genomic signal. The mean Δ*R*^2^ = +0.016 across 51 cell lines, and median *R*^2^ = 0.627 across 601 external cell lines spanning 27 tissue types, provides strong evidence that the genomic image encoder has learned biologically meaningful latent representations that transfer across cancer types.

The tissue-type gradient in external validation — BREAST highest, epithelial cancers intermediate, haematopoietic lowest — is exactly the biological signature expected from an encoder that maps cell lines by genomic content rather than tissue-type label. Haematopoietic malignancies have a fundamentally different genomic landscape from solid epithelial tumours, and the degraded performance on this tissue type is a sign of biological honesty, not a failure of the model.

### 5.3 GradCAM Recovery of Known Cancer Drivers as Biological Validation

The recovery of TP53 among the top-five cross-cell-line genomic activators (5/51 cell lines) – from drug response supervision alone, without pathway annotation — provides partial, rather than dominant, biological validation of the learned representation. TP53 is the most frequently mutated tumour suppressor in breast cancer, with loss-of-function mutations present across diverse subtypes and known to modulate drug sensitivity through multiple mechanisms including apoptotic priming and cell cycle checkpoint abrogation, and its appearance near the top of the frequency ranking is consistent with this central pharmacological role. However, at this checkpoint TP53 does not clearly separate from the surrounding uncharacterised candidates: the gene ranking marginally above it (FSIP2, 6/51) has no current annotation as a breast cancer driver, and BRCA2 — also a curated driver — appears further down the list (3/51). We did not identify PIK3CA among the most frequently activated genes in the aggregate ranking at this checkpoint, despite its high mutation prevalence in breast cancer; this is consistent with PIK3CA’s known mutation-specific, cell-line-restricted pharmacological role (Section 4, single-cell-line case studies), which would not be expected to produce a high frequency count across an aggregate of 51 cell lines, but it also means the present checkpoint does not, on its own, provide an unambiguous signal that the encoder has specifically isolated PIK3CA’s context-dependent role.

Taken together with the BRCA2 and TP53 single-cell-line case studies (Section 4, Figures 8-9), the present checkpoint indicates that the genomic encoder recovers known cancer drivers as part of, rather than as the clearly dominant component of, its learned genomic attribution signal. We consider this a more conservative and more honestly reported characterisation of the model’s interpretability properties than would be obtained by reporting only the most favourable training checkpoint, and we discuss the checkpoint-dependence of this result further below.

GradCAM attribution patterns were broadly consistent between early training checkpoints (epoch 12, val *R*^2^ = 0.680, and epoch 29, val *R*^2^ = 0.715), with TP53 as the dominant cross-cell-line activator in both. However, at the best-validation-*R*^2^ checkpoint reached later in training (epoch 52, val *R*^2^ = 0.718; Section 4), TP53’s relative ranking weakens and an uncharacterised gene (FSIP2) becomes marginally more frequent. This indicates that the driver-gene-frequency signal is not fully stable across training and should be interpreted as checkpoint-dependent rather than as a fixed property of the architecture; we report the later, best-validation checkpoint throughout this revision in the interest of using the single most defensible model selection criterion (validation *R*^2^) rather than selecting whichever checkpoint yields the most favourable interpretability narrative. The continued presence of TP53 and BRCA2 among the top-ranked genes at both checkpoints, alongside a substantial and only partially overlapping set of uncharacterised candidates, suggests that the encoder has learned a genomically grounded but not yet fully disentangled representation, in which curated drivers are recovered without pathway supervision but do not yet cleanly separate from co-varying uncharacterised genes. We treat this as a genuine limitation of the current encoder (Section 5.4) rather than as a fully resolved validation result.

### 5.4 Limitations and Future Directions

#### Training data scope

DL4DR is trained exclusively on breast cancer cell lines. While external validation demonstrates cross-cancer transfer, the genomic encoder’s latent space is anchored by breast cancer biology. Extending training to the full DepMap pan-cancer dataset (1,379 cell lines) would densify the latent space and improve coverage for genomically distant cancer types (Path A). We note separately that the leave-cell-line-out result, while validated to be free of identifier- and image-level data leakage (Section 3.4), is nonetheless estimated from only 51 breast cancer cell lines; the relatively constrained genomic and molecular subtype diversity within a single-tissue pool of this size may itself inflate apparent generalisation relative to what would be observed across a larger, pan-cancer cell-line pool, independent of any leakage concern. This distinction - real generalisation within a narrow biological neighbourhood versus generalisation across more genomically distant contexts — is precisely what the pan-cancer extension in Path A is designed to test.

#### Genomic encoder architecture

The current CNN encoder operates on pixel-level genomic images without knowledge of gene regulatory networks or pathway structure. Replacing it with a pre-trained genomic foundation model (Geneformer [Theodoris et al., 2023], scGPT [Cui et al., 2024]) while keeping the compound tower and fusion layer unchanged would leverage cross-tissue pre-training and likely improve both performance and biological interpretability (Path B).

#### Compound coverage

The 59% of compounds tested in only one cell line limits learning of drug-cell-line interaction effects. Focused experimental designs that test a panel of structurally diverse compounds across all cell lines would substantially improve interaction signal.

#### Checkpoint-dependence of interpretability results

As noted in Section 5, the cross-cell-line driver-gene-frequency ranking is not fixed across training: at the best-validation-*R*^2^ checkpoint (epoch 52) reported throughout this revision, TP53 is recovered among the top-five most frequently activated genes but is not clearly dominant over uncharacterised candidates, whereas an earlier checkpoint (epoch 29) had shown a more pronounced TP53 signal. Predictive performance (val *R*^2^) and interpretability-signal clarity are therefore not guaranteed to improve in lockstep over training, and a single checkpoint’s GradCAM ranking should not be over-interpreted as a stable, architecture-level property. A more rigorous treatment would report driver-gene recovery aggregated or averaged across multiple checkpoints, or as a function of training epoch, rather than from a single snapshot; we leave this checkpoint-robustness analysis to future work.

#### Agentic retrieval-augmented interpretation

The GradCAM outputs presented in Sections 4 and 5 – atom-level attributions for individual compounds (Figures 4–6) and gene-level driver scores for individual cell lines (Figures 8–9) – are numerical outputs that currently require manual literature lookup to contextualise biologically. A natural extension is to attach an LLM agent downstream of these fixed, already-computed attributions: given a ranked list of attributed atoms or genes, the agent would retrieve relevant pharmacological or pathway literature and compose a natural-language rationale linking the model’s numerical attribution to known mechanism. We emphasise that the role of the agent here is strictly interpretive and retrieval-based, not representational: the agent would not replace the D-MPNN, ECFP, or CNN encoders, nor generate or modify the SMILES or genomic-image inputs themselves, since natural-language descriptions of molecular or genomic structure are intrinsically lossier and more hallucination-prone than the structured representations already used by DL4DR. Agentic orchestration is treated here as a means of making existing quantitative outputs more interpretable to domain experts, not as an end in itself or a substitute for the model’s structural representations (Path C).

#### Specific Aim 2: Perturbation-aware modeling

Beyond static genomic backgrounds, drug activity is strongly modulated by biological perturbations such as CRISPR knockouts, pathway activation states, epigenetic alterations, drug priming, and transcriptomic shifts induced by chemical or genetic interventions (e.g., L1000/CMap signatures). These perturbations reshape target abundance, pathway wiring, compensatory mechanisms, and synthetic-lethal interactions, thereby altering drug sensitivity. A natural extension of DL4DR is to incorporate perturbations as a third modality – alongside compound structure and baseline genomics – via a perturbation encoder or conditional modulation layers (e.g., FiLM, conditional attention). This would enable the model to predict context-specific drug responses under defined interventions and to simulate counterfactual scenarios such as “How would IC50 change if gene X were knocked out?’. Integrating perturbation-aware modeling would move DL4DR toward a more mechanistic, intervention-capable framework aligned with precision oncology (Path D).

## 6 Conclusion

Ve presented DL4DR, a Two-Tower Residual Late Fusion model that addresses two fundamental limitations of existing drug response prediction approaches: the over-smoothing trap in pure graph neural networks, and the inability of lookup-table cell line encoders to generalise to unseen biological contexts. By combining ECFP hard memorization with D-MPNN and ORNN soft representations in an additive residual framework, and encoding cell lines as content-based genomic images rather than indexed embeddings, DL4DR achieves consistent improvement over the ECFP-only baseline across 48/51 breast cancer cell lines (94.1%) and transfers to 6 1 external cell lines spanning 27 cancer tissue types (median *R*^2^ = 0.627; statistically significant positive correlation between genomic distance and prediction error, *r* = 0.159, *p* = 9.2 *×* 10^−5^). GradCAM analysis of the Cell Line Tower recovers known cancer drivers including TP53 and BRCA2 among its most frequently activated genes without pathway supervision, although at the best-validation checkpoint reported here these curated drivers do not yet clearly dominate over uncharacterised candidate genes — a checkpoint-dependent limitation we discuss explicitly in Section 5.4 rather than treating as a fully resolved result.

The residual decomposition principle — memori e what is known, learn the deviation — is a broadly applicable design pattern for sparse matrix problems where interaction effects must be distinguished from main effects. We make all code, pretrained checkpoints, and genomic images available at https://github.com/bayjuan5/DL4DR to facilitate extension to pan-cancer settings and integration with genomic foundation models.

## Acknowledgements

This work was supported in part by computational resources provided by the Texas Advanced Computing Center (TACC) under allocation MCB23032. Model training was performed on Google Colaboratory using NVIDIA Tesla T4 GPU resources. The authors thank the DepMap consortium and the Genomics of Drug Sensitivity in Cancer (GDSC) project for making pharmacogenomic datasets publicly available, and Claude (Anthropic) for assistance with L^A^T_E_X manuscript preparation and interpretability visualisation development.

